# A novel glucocorticoid and androgen receptor modulator reduces viral entry and innate immune inflammatory responses in the Syrian Hamster model of SARS-CoV-2

**DOI:** 10.1101/2021.02.20.432110

**Authors:** Savannah M. Rocha, Anna C. Fagre, Amanda S. Latham, Katriana A. Popichak, Casey P. McDermott, Clinton C. Dawson, Jason E. Cummings, Juliette Lewis, Philip Reigan, Tawfik A. Aboellail, Rebekah C. Kading, Tony Schountz, Neil D. Theise, Richard A. Slayden, Ronald B. Tjalkens

## Abstract

Since its initial discovery in late 2019, severe acute respiratory syndrome coronavirus 2 (SARS-CoV-2), the cause of COVID19, has spread worldwide and despite significant research efforts, treatment options remain limited. Replication of SARS-CoV-2 in lung is associated with marked infiltration of macrophages and activation of innate immune inflammatory responses triggered, in part, by heightened production of interleukin-6 (IL-6) that recruits lymphocytes to the site of infection that amplify tissue injury. Antagonists of the glucocorticoid and androgen receptors have shown promise in experimental models of COVID19 and in clinical studies, because cell surface proteins required for viral entry, angiotensin converting enzyme 2 (ACE2) and the transmembrane serine protease 2 (TMPRSS2), are transcriptionally regulated by these receptors. We therefore postulated that the glucocorticoid (GR) and androgen receptor (AR) antagonist, PT150, would reduce infectivity of SARS-CoV-2 and prevent inflammatory lung injury in the Syrian golden hamster model of COVID19. Animals were infected intranasally with 2.5 × 10^4^ TCID50/ml equivalents of SARS-CoV-2 (strain 2019-nCoV/USA-WA1/ 2020) and PT150 was administered by oral gavage at 30 and 100 mg/Kg/day for a total of 7 days. Animals were then examined at days 3, 5 and 7 post-infection (DPI) for lung histopathology, viral load and production of proteins regulating the initiation and progression of SARS-CoV-2 infection. Results of these studies indicated that oral administration of PT150 decreased replication of SARS-CoV-2 in lung, as well as expression of ACE2 and TMPRSS2 protein. Hypercellularity and inflammatory cell infiltration driven by macrophage responses were dramatically decreased in PT150-treated animals, as was tissue damage and expression of IL-6. Molecular modeling suggested that PT150 binds to the co-activator interface of the ligand binding domain of both AR and GR and thereby acts as an allosteric modulator and transcriptional repressor of these receptors. Phylogenetic analysis of AR and GR across multiple species permissive to SARS-CoV-2 infection revealed a high degree of sequence identity maintained across species, including human, suggesting that the mechanism of action and therapeutic efficacy observed in Syrian hamsters would likely be predictive of positive outcomes in patients. PT150 is therefore a strong candidate for further clinical development for the treatment of COVID19 across variants of SARS-CoV-2.

## Introduction

Emerging in late December 2019 in Wuhan, China, several unidentified cases of severe pneumonia were reported of which were epidemiologically linked to a seafood and wet animal wholesale market. Through deep sequencing of lower respiratory samples from these patients, a novel betacoronavirus was identified [1, 2] and has since been distinguished as the initial point source of the biological threat that has now developed into a global pandemic. As of late 2020, severe acute respiratory syndrome coronavirus 2 (SARS-CoV-2) has infected over 40 million people and has been responsible for over 1.14 million deaths [3] and continues to surge in many countries. The infection can be spread through respiratory droplet transmission from asymptomatic, pre-symptomatic, and symptomatic carriers [4] making diagnosis and quarantine efforts difficult, ultimately leading to increased propagation and dissemination of infectious virus.

Coronaviruses (Order: *Nidovirales*; Family: *Coronaviridae*) are enveloped, non-segmented, positive-sense single-stranded RNA viruses that contain very large genomes up to 33.5 kilobases (kb). The four subtypes (genera) of these viruses – alpha, beta, gamma, and delta-coronaviruses – share a highly conserved genome organization comprising a large replicase gene followed by structural and accessory genes. The organization of the coronavirus genome is organized from the 5’-leader-UTR, replicase, S(spike), E(envelope), M(membrane), N(nucleocapsid) to the 3’ UTR poly (A) tail [5]. Notably, production of the spike protein has been linked to severity of disease. The spike protein is a proteolytically processed glycoprotein that extends from the viral membrane and modulates virus-cell membrane fusion. In order to acquire functionality, the protein must undergo multiple stepwise endoproteolytic cleavages [6]. The spike glycoprotein of SARS-CoV-2 has two domains, the S1 domain comprising residues 12 – 667 and the S2 domain, comprising residues 668-1273. The S1 subunit contains the receptor binding domain (RBD), which interacts with the ACE2 receptor, whereas the S2 subunit remains associated with the viral envelope [7]. Viral infection requires proteolytic cleavage at Arg685-Ser686 at the S1 site by the transmembrane protease, serine 2 (TMPRSS2), followed by cleavage at the S2 site at Arg815-Ser816 [6]. Proteolytic cleavage of the spike protein then enables membrane fusion and entry into the host cell in complex with the ACE2 receptor [8].

Downregulating expression of ACE2 and TMPRSS2 has therefore emerged as an important therapeutic strategy for the treatment of COVID19 in order to increase host defense by preventing entry of SARS-CoV-2 into cells, thereby limiting viral replication. Clinical evidence supports this hypothesis, where a prospective study reported a decrease in the rate of intensive care unit admissions in men who had been prescribed anti-androgens for six months prior to hospitalization [9]. Both ACE2 and TMPRSS2 are highly expressed in bronchiolar epithelial cells and transcriptionally regulated at the promoter level through the androgen receptor (AR) [8]. In a model of SARS-CoV-2 infection, inhibitors of AR transcriptional activity markedly reduced viral infectivity and decreased inflammatory lung injury in experimental animals [8]. Both ACE2 and TMPRSS2 are also significantly regulated by inflammation. ACE2, as well as the inflammatory cytokine interleukin-6 (IL-6), are non-canonical interferon-responsive genes (ISGs) that are highly expressed following infection with SARS-CoV-2 [10]. Inflammation also directly upregulates expression of TMPRSS2 [11]. Steroids have therefore been extensively used to treat COVID19 patients, albeit with mixed results. A recent study in *The Lancet* indicated that clinical evidence does not support corticosteroid treatment for SARS-CoV-2-related lung injury. Patients who were given corticosteroids were more likely to require mechanical ventilation, vasopressors, and renal replacement therapy [12]. Corticosteroids such as dexamethasone have been shown to increase expression of ACE2 [13, 14], which would enhance viral entry and replication and could therefore worsen infection when given too early. Dexamethasone also modestly increased expression of TMPRSS2 in these studies, which could likewise potentially enhance membrane fusion and vesicular uptake of SARS-CoV-2. Neither hydrocortisone nor prednisolone were able to decrease expression of TMPRSS2 [13]. Thus, better therapeutics are needed that can modulate these receptor systems to downregulate expression of proteins critical to viral entry.

During viral RNA synthesis, both genomic and sub-genomic RNAs are produced that serve as mRNAs for the structural and accessory proteins that are downstream of viral replicase proteins [5]. SARS-CoV-2 contains up to seven stem-loop regulatory sequences in the 5’-UTR that that are thought to control stages of RNA synthesis and may confer species specificity for virulence [5, 15]. In part because a number of human RNA viruses have glucocorticoid receptor *cis*-acting elements in their 5-UTR’s, antagonists of the glucocorticoid receptor are one potential class of drugs for therapeutic modulation of SARS-CoV-2 infection. A report comparing the host inflammatory response to SARS-CoV (SARS1) and HCoV-EMC (MERS) revealed a unique set of 207 genes dysregulated during the course of infection. Based on these data, the authors predicted that selected kinase inhibitors and glucocorticoid receptor modulators could function as potential antiviral compounds [16]. These studies suggested two important points about modulating infection with human coronaviruses: 1) targeting cellular responses has been shown to inhibit viral replication and 2) immunomodulatory drugs that reduce the excessive host inflammatory response to respiratory viruses have therapeutic benefit, as seen with influenza virus infections [16].

However, classical antagonists of glucocorticoid function that compete for interaction at the steroid binding pocket of the receptor ligand binding domain (LBD) could be problematic, due to excessive blockade of cortisol function. Thus, allosteric modulators of both AR and GR that could dampen transcriptional activation through these receptors would be preferable. Such interactions tend to favor stabilization of transcriptional co-repressor proteins on chromatin, such as CoREST, HDAC2/3/4 and NCoR2, that prevent binding of co-activator proteins in response to activation of *cis*-acting transcription factors [17, 18]. To address the potential for a transcriptional modulator to protect against SARS-CoV-2 infection through downregulation of AR/GR-dependent expression of ACE2 and TMPRS22, we examined the therapeutic efficacy of (11β,17β)-11-(1,3-benzodioxol-5-yl)-17-hydroxy-17-(1-propynyl)-estra-4,9-dien-3-one (designated as “PT150”), a synthetic AR and GR modulator shown to antagonize human GR [19, 20]. In a screening study conducted by the National Institute of Allergy and Infectious Diseases, PT150 had broad inhibitory activity towards several RNA viruses, including influenza, Zika virus and β-coronaviruses, and was recently shown to have direct anti-viral effects against SARS-CoV-2 in human bronchiolar epithelial cells *in vitro* [21]. Based on these data, and on previous host-pathogen interaction studies, we postulated that PT150 would be an effective inhibitor of SARS-CoV-2 infection *in vivo* in the Syrian golden hamster model of COVID19. The results of this study demonstrate that PT150 given orally once daily for 7 days prevented replication of SARS-CoV-2 in lung, decreased infiltration of macrophages, improved lung pathology and reduced expression of both ACE2 and TMPRSS2.

## Materials and Methods

### Virus and cell culture procedures

SARS-CoV-2 strain 2019-nCoV/USA-WA1/2020 was obtained from BEI Resources and propagated on Vero cells (ATCC CCL-81, American Type Culture Collection) at 37°C with 5% CO2. Virus titrations from animal tissues were performed by plaque assay as described previously [21]. Plaques were visualized two days post inoculation following fixation with 10% formalin for 30 minutes, and addition of 0.25% crystal violet solution in 20% ethanol.

### Animal procedures

All animal protocols were approved by the Institutional Animal Use Committee at. Colorado State University (IACUC Protocol No. 996), hamsters were handled in compliance with the PHS Policy and Guide for the Care and Use of Laboratory Animals and procedures were performed in accordance with National Institutes of Health guidelines. Male and female Syrian hamsters were used in studies at 8 weeks of age (N=60, Charles River Laboratory). The animals were housed in the CSU animal facility and allowed access to standard pelleted feed and water *ad libitum* prior to being moved to the Biosafety Level 3 containment facility for experimental infection. Hamsters were anesthetized by inhalation with isoflurane and then intranasally inoculated with 2.5 × 10^4^ TCID50/ml equivalents of SARS-CoV-2 (strain 2019-nCoV/USA-WA1/ 2020) in sterile Dulbecco’s Modified Eagles Medium (DMEM). Hamsters not receiving SARS-CoV-2 were given a sham inoculation with the equivalent volume of DMEM vehicle. To assess activity of PT150 (supplied by Palisades Therapeutics/Pop Test Oncology LLC) against SARS-CoV-2, experimental groups (*N*=6 animals per group at each timepoint) were as follows: control (sham inoculation + miglyol vehicle), SARS-CoV-2 + miglyol, SARS-CoV-2 + 30 mg/Kg PT150, SARS-CoV-2 + 100 mg/Kg PT150. The experimental drug (PT150) was dissolved in 100% miglyol 812 and delivered by oral gavage at 8 μL/gm body weight under isoflurane anesthesia. Animals were weighed daily to deliver an accurate dose of drug. The SARS-CoV-2 + vehicle group also received miglyol 812 by oral gavage. Animals were observed for clinical symptomology at time of dosing each day (lethargy, ruffled fur, hunched back posture, nasolacrimal discharge, and rapid breathing). Groups of animals were terminated at 3, 5 and 7 days post-infection (DPI). Eighteen hamsters (6 SARS-CoV-2 + vehicle, 6 SARS-CoV-2 + 30 mg/Kg PT150; 6 SARS-CoV-2 + 100mg/Kg PT150) were euthanized at 3 and 5 DPI. On day 7 post-infection (7 DPI) the remaining 24 hamsters were euthanized (6 DMEM + miglyol; 6 SARS-CoV-2 + vehicle, 6 SARS-CoV-2 + 30 mg/Kg PT150; 6 SARS-CoV-2 + 100mg/Kg PT150). Animals were terminated by decapitation under isoflurane anesthesia and tissue was collected for immunohistochemistry, viral isolation, RNA analysis and histopathology.

### Histopathology

Lungs from 60 hamsters were extirpated *en bloc* and fixed whole in 10% neutral buffered formalin under Biosafety Level 3 (BSL-3) containment for at least 72 hours before being transferred to CSU Veterinary Diagnostic Laboratory, BSL-2 necropsy area for tissue trimming and sectioning. Four transverse whole-lung sections were stained with hematoxylin and eosin (H&E). Tissue was sectioned at 5μm thickness and were mounted onto poly-ionic slides. Sections were then deparaffinized and immunostained using the Leica RX^m^ automated robotic staining system. Antigen retrieval was performed by using Bond Epitope Retrieval Solutions 1 and 2 for 20 minutes each in conjunction with base plate heat application. Sections were then permeabilized (0.1% Triton X in 1X TBS) and blocked with 1% donkey serum. Primary antibodies were diluted to their optimized dilutions in tris-buffered saline and incubated on the tissue for 1 hour/antibody: Rabbit SARS nucleocapsid protein (SARS-CoV-2; Rockland; 1:500), goat ionized calcium binding adaptor molecule 1 (IBA1; Abcam; 1:50), goat angiotensin converting enzyme 2 (ACE2;R&D Systems; 1:500), rabbit transmembrane serine protease 2 (TMPRSS2; Abcam; 1:500), mouse interleukin 6 (IL-6; ThermoFisher; 1:500). Sections were then stained for DAPI (Sigma) and were mounted on glass coverslips using ProLong Gold Anti-Fade medium and stored at 4°C until imaging.

### Immunofluorescence Imaging and Protein Quantification

The studies described here were conducted by a single investigator. Images were captured using an automated stage Olympus BX63 fluorescent microscope equipped with a Hamamatsu ORCA-flash 4.0 LT CCD camera and collected using Olympus CellSens software. Quantification of protein was performed by acquiring five randomized images encompassing the pseudostratified columnar epithelium around bronchi at 400x magnification (Olympus X-Apochromat air objective; N.A. 0.95) all from different lung lobes. Regions of interest were then drawn to enclose the epithelial layer and exclude the lumen, in order to accurately obtain average intensity measurements. The Count and Measure function on Olympus CellSens software was then used to threshold the entirety of the ROI and measure the given channel signal. Quantification of invading inflammatory cells was performed by generating whole lung montages by compiling 100x images acquired using automated stage coordinate mapping with an Olympus 10X air objective (0.40 N.A.) All images were obtained and analyzed under the same conditions for magnification, exposure time, lamp intensity, camera gain, and filter application. ROIs were drawn around the lung sections and the co-localization function of Count and Measure within Olympus CellSens software was applied the sections. IBA1+ cells were determined per 1mm^2^ areas given the previously drawn ROI overall area.

### Quantification of Inflammatory Cell Infiltration and Consolidation

Quantification of the total affected pulmonary parenchyma as well as counting of inflammatory cells per area (region of interest, ROI, 1mm^2^) was determined in hematoxylin and eosin (H&E) - stained histological sections by digital image analysis. A digital montage was compiled at 100X magnification using an Olympus X-Apochromat 10X air objective (N.A. 0.40) consisting of approximately 1,200 individual frames per lung lobe. Affected regions of interest (ROI’s) were subsequently automatically identified using Olympus CellSens software by quantifying whole-lung montages scanned from each hamster for total number of nuclei or nucleated cells (to exclude erythrocytes) stained with H&E, relative to the total area of the ROI for each lung.

### Computational-based Modeling

PT150 was docked into the crystal structures of the ligand-binding domain of the androgen receptor (PDB: 2PIT) [22], and the glucocorticoid receptor (PDB: 3CLD) [23], using the Glide module within Schrödinger (Release 2020-2, Schrödinger LLC, New York, NY) [24–26]. Prior to docking, the water molecules were removed, and the proteins were prepared by assigning bond orders, adding hydrogens, and repairing any side chains or missing amino acid sequences. To complete protein preparation a restrained minimization of the protein structure was performed using the default constraint of 0.30Å RMSD and the OPLS_2005 force field [27]. The prepared proteins were subjected to SiteMap analysis [26], that identified the available binding sites in the ligand binding domains of the androgen and glucocorticoid receptors and docking grids were generated using Receptor Grid Generation. PT150 prepared using LigPrep by generating possible states at the target pH 7.0 using Epik and minimized by applying the OPLS_2005 force field [27]. Molecular docking simulations were performed targeting each potential binding site for PT150 using the Glide ligand docking module in XP (extra precision) mode and included post-docking minimization [25].

### Phylogenetic Analysis and Similarity Score Representation

Phylogenetic analysis was performed by investigating protein coding sequences across multiple species within the protein of interest. FASTA files were downloaded from National Center for Biotechnology Institute’s (NCBI) gene databases and were then input into Molecular Evolutionary Genetic Analysis (MEGAX, v.10.1.8) software for alignment. Muscle alignment was performed, and a neighbor-joining phylogenetic tree was constructed. Similarity scores between species were generated by utilizing the Basic Local Alignment Search Tool for protein-protein comparison on the National Center for Biotechnology Information interface.

## Results

### Clinical observations and levels of SARS-CoV-2 in the lungs of Syrian hamsters treated with PT150

Oral gavage with PT150 or vehicle began on the same day as infection with 2.5 × 10^4^ TCID_50_ SARS-CoV-2 (USA-WA1-2020 strain) by intranasal inoculation. Body weights were monitored daily for each animal, with a noted decline in average body weight in each experimental group that reached a maximum loss by day 5 with a total overall loss of eight-percent body weight (**Fig 1A,B)**. This finding is consistent with other longitudinal studies in Syrian golden hamsters infected with SARS-CoV-2 that demonstrate the maximal clinical severity of disease at day 5 post-infection [28, 29]. Infected hamsters treated with vehicle-only showed the greatest decline in body weight relative to controls, that was prevented by treatment with PT150 at 30 and 100 mg/Kg/day **(Fig 1A)**. Two-way ANOVA analysis indicated a difference with treatment (*p*<0.0007, F (3,92)=6.231) and time post-infection (*p*<0.0001, F (7,92)=16.39). Hamsters treated with PT150 at 30 and 100 mg/Kg/day did not show a statistically significant difference in body weight from control hamsters at day 5 post-infection, at which the maximal extent of body weight loss is observed in Syrian hamsters infected with SARS-CoV-2 and treated with vehicle only (**Fig 1B**). Hamsters challenged with SARS-CoV-2 showed marked lethargy, ataxia, ruffled fur, altered posture and overall decreased movement and exploratory behavior, clinical signs and behaviors that were observably less pronounced in hamsters treated with 30 mg/Kg/day PT150. Infected hamsters treated with 100 mg/Kg/day PT150 displayed clinical behavior largely indistinguishable from uninfected control animals.

**Figure 1.**
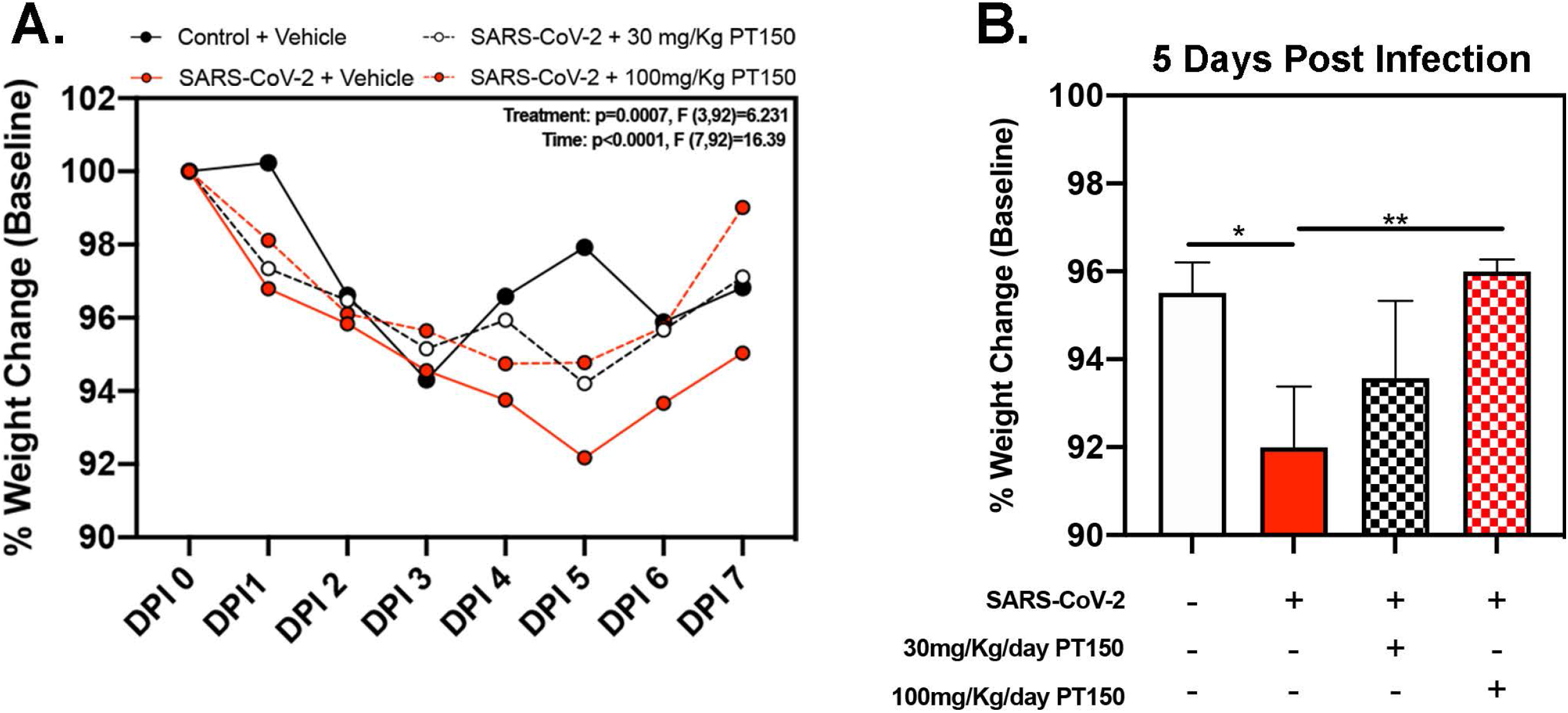
Clinical observations in Syrian hamsters infected with SARS-CoV-2. **A**) Time course of body weights for all groups plotted as percent change from baseline for each animal. Two-way ANOVA analysis indicated significance for both Time (p<0.0001, F (7, 92) = 16.39) and Treatment condition (p=0.0007, F *3, 92) = 6.231). Maximal loss of body weight was observed by five days post-infection. **B**) Body weights were directly compared between groups at 5 days post-infection. Significant differences were detected between the control and SARS-CoV-2 groups, as well as between SARS-CoV-2 + 30 mg/Kg PT150. The SARS-CoV-2 + 100 mg/Kg PT150 group was not different from the SARS-CoV-2 + vehicle group, but was also not different from control. Groups were compared by one-way ANOVA using Tukey’s post hoc test. *p<0.05, **p<0.01. N=6 animals per group.

### Lung histology

Paraffin-embedded lung sections were stained with hematoxylin and eosin (H&E) and examined by a veterinary pathologist blinded to the treatment groups. Representative lung sections from each experimental group are presented in **Figure 2.** In hamsters infected with SARS-CoV-2, affected portions of lungs showed marked histiocytic/neutrophilic broncho-interstitial pneumonia with hyperplasia of pneumocyte type II and formation of both bronchiolar and alveolar syncytial cells (**Fig 2A-C**, low power images and high magnification insets). Inflammation equally involved main branches of the pulmonary artery, where infiltrating macrophages were observed dissecting the tunica media and lifting the tunica intima, with clustering of circulating monocytes along hypertrophied, apoptotic and occasionally hyperplastic endothelial lining cells. Lungs from vehicle-treated infected hamsters show approximately 70% destruction of pulmonary parenchyma by an intense mixed inflammatory infiltrate with marked expansion of alveolar interstitium, as well as peri-bronchial inflammation and arteritis of large pulmonary vessels. Of note, peri-bronchial inflammation and arteritis of a large pulmonary vessel was observed (**Fig 2B**). High power images (**Fig 2B,C**, insets) of the bronchiolar wall show inflammatory cells comprising macrophages and neutrophils dissecting through bronchiolar muscular wall with similar infiltration of several rows of monocytes/macrophages lifting the tunica intima and dissecting the tunica media. By 7 DPI, infected animals treated with vehicle-only (**Fig 2C**) had less inflammation, with clearing of luminal infiltrates in smaller bronchioles. High power images show reduced clustering of monocytes/macrophages in pulmonary arteries with lingering inflammation in medium-sized bronchioles and parenchymal consolidation. Treatment with PT150 at 30 mg/Kg/day (**Fig 2D-F)** and 100 mg/Kg/day (**Fig 2G-I**) alleviated the extent of inflammation in main stem bronchi and respiratory bronchioles, as well as in medium-sized pulmonary arteries. The percentage of normal to affected parenchyma showing significant resolution was markedly increased in PT150-treated animals, particularly in the 100 mg/Kg/day group, even at the peak of inflammatory infiltration by 5 DPI (**Fig 2H**, low and high-power images). In the 100 mg/Kg/day treatment group, there was near complete resolution within the parenchyma by 7 DPI (**Fig 2I**, low and high-power images), relative to untreated animals infected with SARS-CoV-2, which showed significant consolidation and loss of parenchymal structure by 7 DPI (**Fig 2C**, low and high-power images). In addition, the number of apoptotic endothelial cells in pulmonary bronchi was greatly reduced by treatment with 100 mg/Kg/day PT150 (**Fig 2G-I**, high magnification insets). In hamsters treated with 100 mg/Kg/day PT150, there was a marked reduction in the inflammatory response within bronchioles, interstitium and arteries, with almost complete resolution of bronchiolar inflammation and reduced clustering of circulating monocytes to a marginating single row. No medial dissection is observed in these vessels. The overall paradigm for treatment and isolation of tissues for analysis of viral load, gene expression and histopathology is depicted in the diagram in **Fig 2J**.

**Figure 2.**
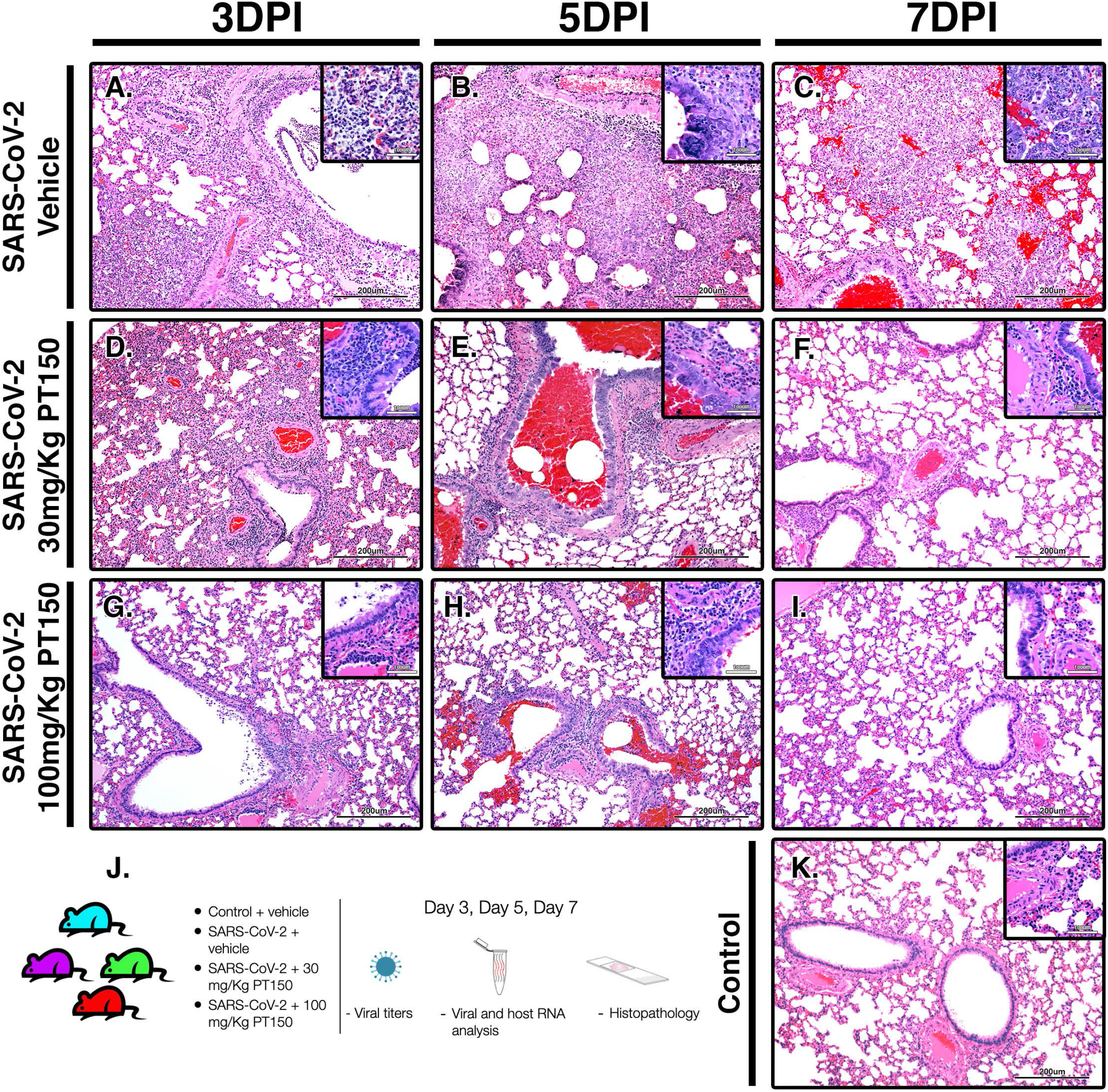
PT150 protects against adverse histopathological outcomes in the lungs of Syrian hamsters infected with SARS-CoV-2. Hamsters were infected by intranasal inoculation with 2.5 × 10^4^ TCID50/ml equivalents of SARS-CoV-2 (strain 2019-nCoV/USA-WA1/ 2020) and lung tissue was stained with hematoxylin and eosin (H&E) for examination of pathological changes on days 3, 5 and 7 post-infection. Treatment groups were (**A-C**) SARS-CoV-2 + vehicle, (**D-F**), SARS-CoV-2 + 30 mg/Kg/day PT150, (**G-I**) SARS-CoV-2 + 100 mg/Kg/day PT150 and (**J**) Control + vehicle. *N*=6 animals per group. Images were collected at 100X overall magnification (large image panels) or 400X overall magnification (inset panels). (**K**) Study timeline and schematic for treatments and analysis of viral titers, host and viral RNA and histopathology.

### Analysis of immune cell infiltration and broncho-interstitial pneumonia in lung tissue of animals exposed to SARS-CoV-2

Lungs were examined for the extent of immune cell infiltration at 3, 5 and 7 days post-infection (DPI) by quantitative digital image analysis (**Fig 3**). Whole mount sections of paraffin-embedded lung tissue were stained with H&E and bright field grayscale images were collected using a microscope equipped with a scanning motorized stage. Pseudo-colored H&E images are depicted in blue, overlaid with ROIs detected by intensity thresholding in red. Hematoxylin-positive immune cell soma were rendered as focal points within the regions of interest to calculate the percent hypercellularity of tissue following infection with SARS-CoV-2. By 3 DPI, lung tissue showed significant infiltration of immune cells in the SARS-CoV-2 + vehicle group in addition to widespread hemorrhaging (**Fig 3A**). Immune cell infiltration was decreased in dose-dependent fashion by treatment with PT150 at 30 and 100 mg/Kg/day (**Fig 3D-F, G-I)**. Uninfected control hamsters treated with vehicle-only, displayed minimal levels of macrophage hypercellularity with clear bronchi and open parenchyma and an absence of inflammatory infiltrate (**Fig 3J**). The percent of total lung area displaying immune cell hypercellularity was quantified by ROI thresholding and normalizing to the total lung section area (**Fig 3K).** This effectively showed the post-infection response mediated by immune-cell infiltration. There was a time-dependent increase in cellular reactivity, tissue pathology and bronchio-interstitial pneumonia. The peak of cellular infiltration occurred at the 7-day timepoint revealing progressive consolidation. These affects are decreases with the administration of PT150 at the 30mg/Kg/day dose as well as the 100mg/Kg/day dose.

**Figure 3.**
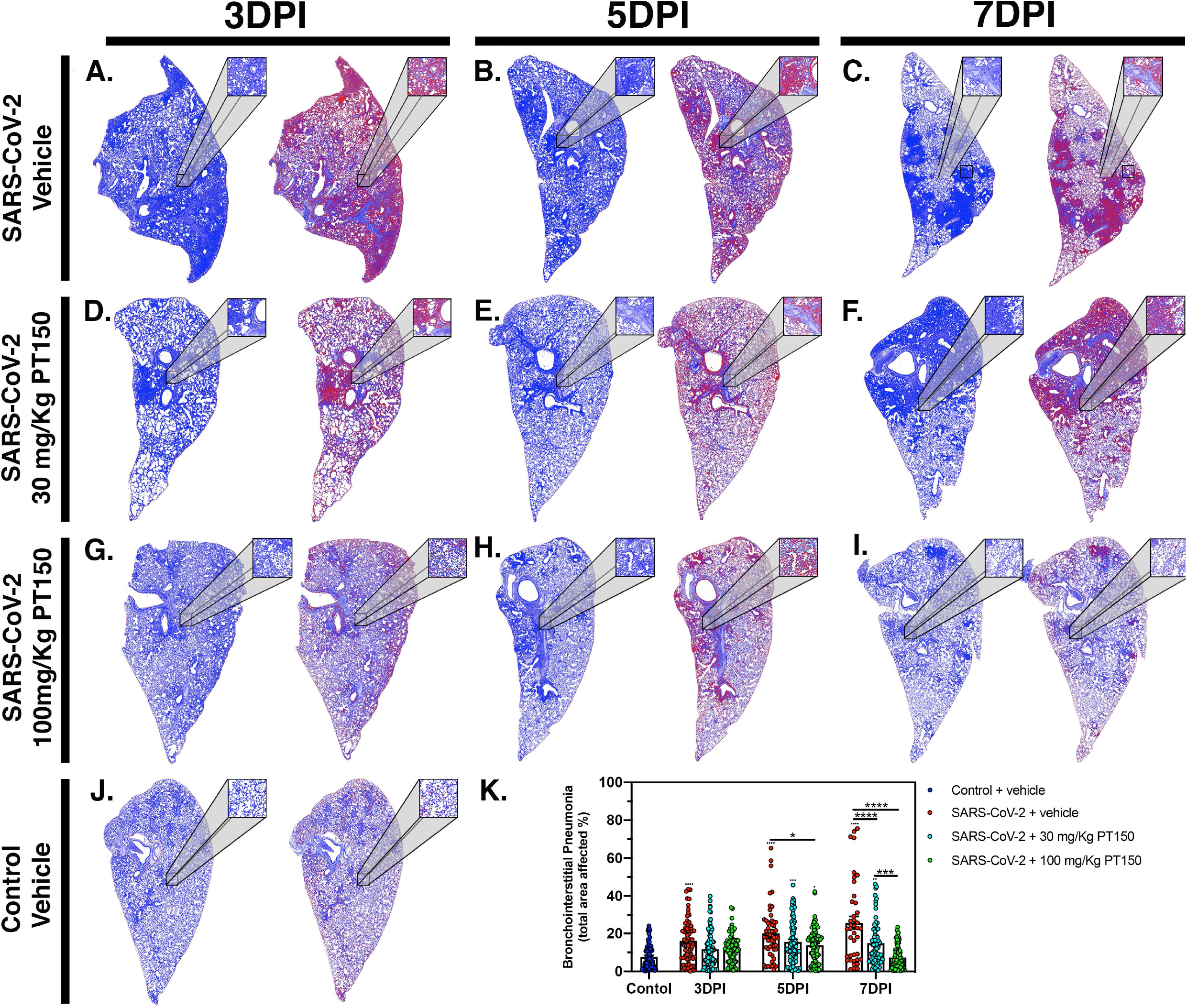
Reduction of immune cell infiltration and broncho-interstitial pneumonia by PT150 treatment. Overall hypercellularity within lung tissue, resulting in pathological broncho-interstitial pneumonia, was determined using ROI delineations on hematoxylin and eosin-stained sections for all groups: days 3, 5 and 7 post-infection. **A-C,** SARS-CoV-2 + vehicle; **D-F**, SARS-CoV-2 + 30 mg/Kg/day PT150; **G-I**, SARS-CoV-2 + 100 mg/Kg/day PT150; B, Control + vehicle. **K**, Quantification of the total area of the lung tissue affected with broncho-interstitial pneumonia was conducted using automated focal point determination within ROIs following manual thresholding. Pseudo colored grayscale images of H&E sections are depicted in blue, overlaid with ROI’s detected by intensity thresholding in red. (\*\*\*\**p*<0.0001; *N*=6 hamsters/group)

### Phylogenetic analysis and molecular docking with the androgen and glucocorticoid receptors

Because expression of TMPRSS2 and ACE2R are regulated through the andro-corticosteroid signaling pathway [7, 11, 30], we examined the genetic sequence identity of the glucocorticoid and androgen receptors across multiple species associated with propagation of SARS-CoV-2 (**Fig 4**). Comparing sequences of the androgen receptor (**Fig 4A**) indicated a high degree of identity between all species analyzed, with sequence concordance values with the human gene ranging from 84.62 (golden hamster) and 85.47 (greater horseshoe bat) to 98.26 (sunda pangolin). Similarly, sequence concordance values with the human gene for the glucocorticoid receptor (**Fig 4B**) ranged from 89.96 (golden hamster) and 90.89 (greater horseshoe bat) to 95.00 (sunda pangolin). Sequence similarity between these genes amongst difference species is relevant for both potential zoonotic propagation of SARS-CoV-2 as well as to the testing of potential therapeutic compounds acting through the androgen and glucocorticoid receptors.

**Figure 4.**
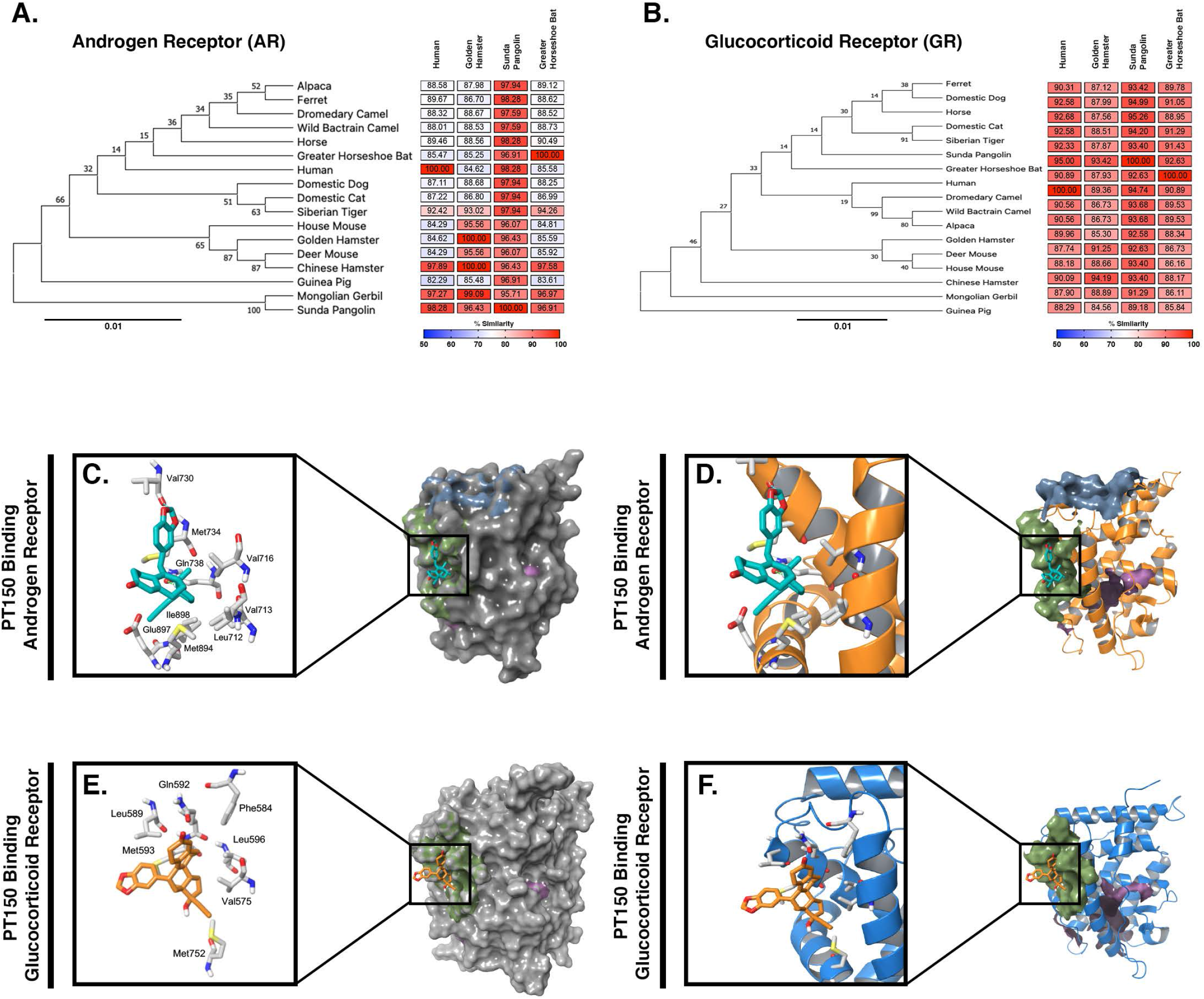
Phylogenetic analysis and molecular docking of PT150 with the androgen and glucocorticoid receptor. Phylogenetic analysis of the androgen receptor (**A**) and glucocorticoid receptor (**B**) indicate a high degree of homology across species, including between human and hamster, indicating that the hamster is an appropriate predictive model to test the protective effects of PT150 against SARS-CoV-2 via modulation of these receptors. The putative binding site on the androgen (**C**) and glucocorticoid (**D**) receptors was analyzed by computer modeling studies using the human receptor.

PT150 was docked into the crystal structures of the ligand-binding domain of the androgen receptor (PDB: 2PIT) and the glucocorticoid receptor (PDB: 3CLD), using the Glide module within Schrödinger (Release 2020-2, Schrödinger LLC, New York, NY) [9–11]. For the androgen receptor (**Fig 4C,D**), the PDB: 2PIT structure was selected due to the 1.76Å resolution and good coverage of the ligand binding domain of the androgen receptor (251 residues). PT150 showed optimal binding to the peptide activator allosteric site on the androgen receptor but did not dock into the steroid binding site identified for dihydrotestosterone. For the glucocorticoid receptor, the PDB: 3CLD was selected due to the 2.84Å resolution, limited other available structures and good coverage of the ligand binding domain of the glucocorticoid receptor (259 residues). Similar to the analysis of the androgen receptor, PT150 preferentially bound to the co-activator peptide allosteric binding site of the glucocorticoid receptor (**Fig 4E,F**). No anchoring interactions were observed for the steroid binding pocket.

### ACE2 and TMPRSS2 production and regulation in animals infected with SARS-CoV-2

The ACE2 receptor and the cell surface TMPRSS2 serine protease are required for viral binding and S-protein priming to facilitate entry of SARS-CoV-2 into cells. We therefore examined production of these proteins in lung tissue from infected hamsters with and without PT150 treatment. ACE2 intensity measurements were quantified by immunofluorescence imaging (**Fig 5**), where pseudostratified columnar epithelium lining bronchioles showed marked decreases in ACE2 production relative to vehicle control (**Fig 5J**) with the administration of 100mg/kg/day of PT150 at the 3-day timepoint as well as the 7-day timepoint (**Fig 5G-I, K**). Decreased ACE2 production was also observed in the 30mg/kg/day PT150 lung sections at 7DPI (**Fig 5F**). Production of ACE2 in vehicle-treated SARS-CoV-2 lung sections were similar to control until the 7-day timepoint, at which point production reached a maximum (**Fig 5A-C**). Expression of TMPRSS2 within bronchiolar cells was increased in the untreated lung sections infected with SARS-CoV-2 (**Fig 6A-C, K**), indicating induction associated with enhanced levels of viral entry, replication and dissemination. In contrast, the 100mg/kg/day PT150 treated animals showed significant decreases in TMPRSS2 protein levels at all timepoints (**Fig 6G-I**) similar to levels observed in the control group (**Fig 6J-K**). There was also reduction in TMPRSS2 proteins levels within the 30 mg/kg/day treatment groups at 5DPI and 7DPI (**Fig 6D-F, K**), similar to levels in control animals at 7DPI.

**Figure 5.**
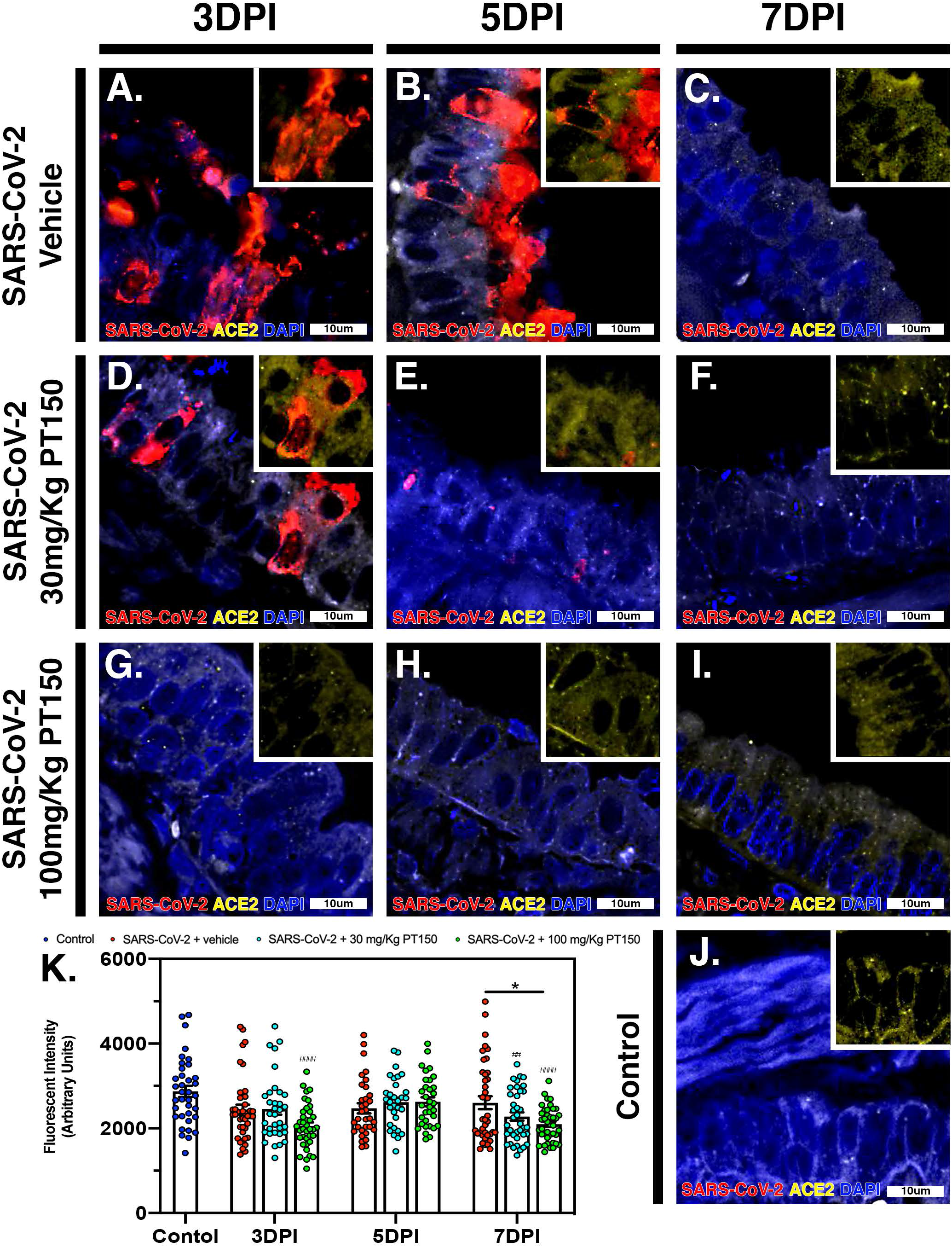
PT150 treatment decreases expression of the ACE2 receptor in lung. In Syrian golden hamsters infected with SARS-CoV-2, there was a modest decrease in expression of the ACE2R in bronchiolar epithelial cells at day 7 post-infection in animals given PT150 at 100 mg/Kg/day. Experimental groups were (A-C) SARS-CoV-2 + vehicle, (D-F), SARS-CoV-2 + 30 mg/Kg/day PT150, (G-I) SARS-CoV-2 + 100 mg/Kg/day PT150, (J) Control + vehicle. K, Quantification of ACE2R expression in lung tissue. *p<0.05

**Figure 6.**
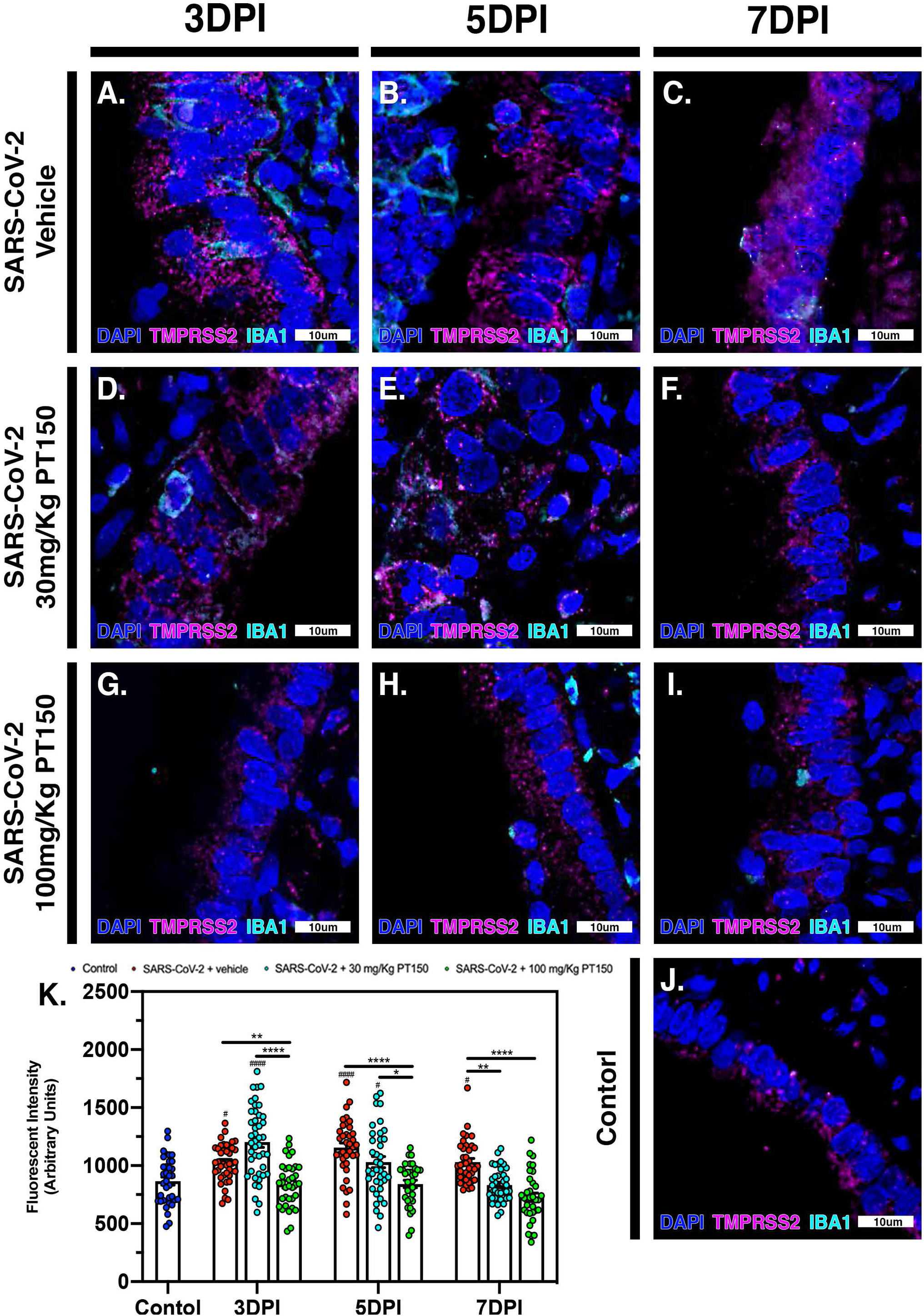
PT150 administration decreases overall TMPRSS2 expression in bronchiolar epithelial cells. In Syrian golden hamsters infected with SARS-CoV-2, there was a significant decrease in expression of the TMPRSS2 in bronchiolar epithelial cells at all timepoints in animals given PT150 at 30 and 100 mg/Kg/day. Experimental groups were (A-C) SARS-CoV-2 + vehicle, (D-F), SARS-CoV-2 + 30 mg/Kg/day PT150, (G-I) SARS-CoV-2 + 100 mg/Kg/day PT150, (J) Control + vehicle. K, Quantification of TMPRSS2 protein expression in lung tissue. *p<0.05, **p<0.01, ****p<0.0001

### Inflammatory activation of macrophages and release of interleukin-6 is decreased by PT150 treatment in Syrian hamsters infected with SARS-CoV-2 in parallel with decreased viral load

The peak of infectivity and viral replication within the Syrian hamster model at sampled time points was observed at 3DPI. Using quantitative immunofluorescence scanning microscopy, we evaluated the extent of lung area containing both SARS-CoV-2 viral protein and infiltrating IBA1+ macrophages (**Fig 7**). In the SARS-CoV-2 + vehicle group, staining for viral nucleocapsid protein indicated the peak of viral protein production at 3DPI, which declined at both 5 and 7DPI (**Fig 7A-E**). Increased infiltration of macrophages was present at 3DPI, peaked at 5DPI and was then followed by a decline at 7DPI **(Fig 7A-E)**. Control animals (**Fig 7S,T**) showed no staining for SARS-CoV-2 and only background levels of IBA1. Viral replication was decreased by administration of 30mg/kg/day of PT150 (**Fig 7G-L**), and was further decreased by the administration of 100mg/kg/day of PT150 (**Fig 7M-R**). In parallel to the decrease in viral nucleocapsid protein, there was a decrease in the percent of total lung area occupied by infiltrating macrophages at 30 and 100 mg/Kg/day PT150 (**Fig 7G-L,M-R**). Quantification of immunofluorescence staining, as measured by the percent of total sampled lung area positive for SARS-CoV-2, revealed marked decreases in both percent lung area expressing viral replication, as well as the number of IBA1+ cells/mm^2^ (**Fig 7U,V**). This demonstrates a dose-response relationship in therapeutic efficacy of PT150 for reducing the viral burden of SARS-CoV-2 in lung, as well as a corresponding decrease in the extent of infiltrating macrophages.

**Figure 7.**
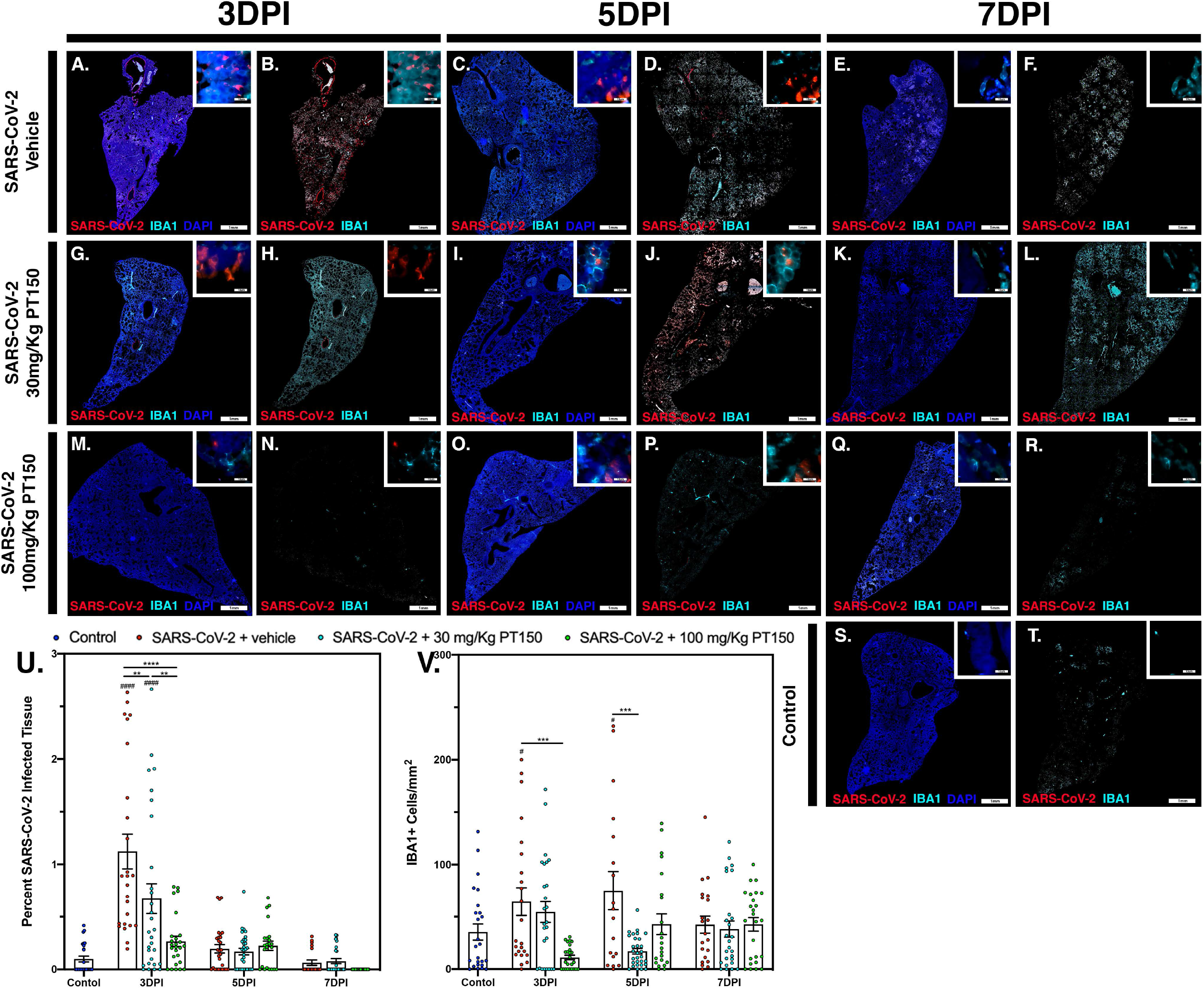
SARS-CoV-2 viral protein expression and macrophage infiltration is reduced by oral administration of PT150. Viral load was determined by immunofluorescence staining of the SARS-CoV-2 nucleocapsid protein. Total viral replication and immune cell infiltration is seen within lung sections at the 3DPI, 5DPI, and 7DPI timepoints. **A-F**, SARS-CoV-2+Vehicle; **G-L,** SARS-CoV-2+30 mg/Kg/day PT150; **M-R**, SARS-CoV-2+100mg/Kg/day PT150; **S-T**, Control + Vehicle. Quantification of percent SARS-CoV-2 infected tissue **(U)** and infiltrating macrophages **(V)** was performed using adaptive intensity thresholding and cellular co-localization of protein expression. N=6 animals per group with a sampling of 5 lobes of tissue per animal. Differences were determined by one-way ANOVA, *p<0.03, **p<0.002, ***p<0.0002, ****p<0.0001. ####p<0.0001 compared to control.

In Syrian golden hamsters infected with SARS-CoV-2, there was a significant increase in macrophage-derived IL-6 within the bronchiolar epithelial layer (**Fig 8A-C**). Immunofluorescence images of infected hamster lung tissue at 3DPI revealed cells within the bronchiolar epithelial layer co-producing high levels of IL-6 (green) with SARS-CoV-2 nucleocapsid protein (red). Nuclei were counterstained with DAPI (blue) and IBA1+ macrophages are shown in cyan. In infected animals, triple label immunofluorescence images show cells staining intensely for IL-6 and co-localizing with expression of SARS-CoV-2 nucleocapsids protein, adjacent to IBA1+ macrophages **(Fig 8A and inset)**. Expression of IL-6 persisted at 5 and 7DPI, even after SARS-CoV-2 nucleocapsid protein was no longer evident (**Fig 8B,C**). Treatment with PT150 and 30 mg/Kg/day (**Fig 8D-F**) and 100 mg/Kg/day (**Fig 8G-I**) dramatically decreased expression of IL-6, concordant with decreases in SARS-CoV-2 and in the presence of IBA1+ macrophages. Quantification of fluorescence data indicated differences between PT150-treated groups and infected + vehicle groups at all timepoints (Treatment, *p*<0.0001, F(3,232)=14.71; Timepoint, *p*<0.0001, F(2,232)=29.99), with the greatest differences evident at 3DPI, where both PT150-treated groups were different from the SARS-CoV-2 + vehicle group, as well as from each other, indicating dose-dependent effects on reduction of IL-6 in the bronchiolar epithelial layer of infected hamsters.

**Figure 8.**
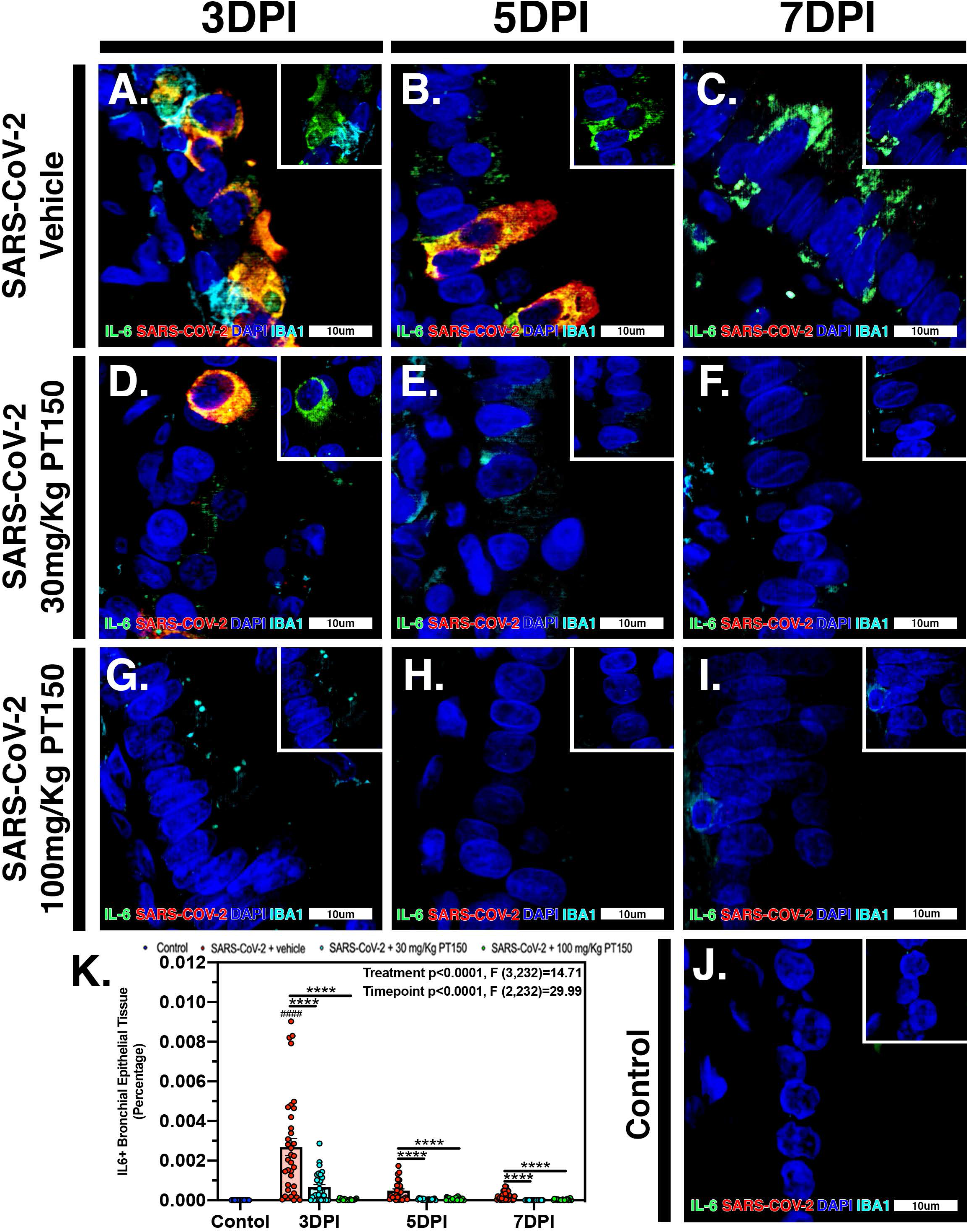
PT150 treatment decreases expression of the inflammatory cytokine Interleukin-6 (IL-6) in lung following infection with SARS-CoV-2. There was a significant decrease in expression of TMPRSS2 in bronchiolar epithelial cells of Syrian golden hamsters infected with SARS-CoV-2 at all timepoints in animals given PT150 at 30 and 100 mg/Kg/day. Experimental groups were (A-C) SARS-CoV-2 + vehicle, (D-F), SARS-CoV-2 + 30 mg/Kg/day PT150, (G-I) SARS-CoV-2 + 100 mg/Kg/day PT150, (J) Control + vehicle. K, Quantification of IL-6 protein expression in lung tissue. \**p*<0.05, \*\**p*<0.01, \*\*\*\**p*<0.0001

## Discussion

Treatment options available for individuals that have SARS-CoV-2 continue to be limited. Of the few treatments that are available, many are the result of repurposing efforts and have low efficacy and limited experimental characterization about the direct mechanisms of action against SARS-CoV-2. Therapeutic approaches have primarily focused on targeting viral proteins [31–33] or the host immune response through corticosteroid administration, which can be detrimental if not administered during the correct period of the infection [34]. However, less research has been published on molecules that have both anti-viral activity as well as immunomodulatory activity to decrease the hyperinflammatory response to SARS-CoV-2 infection. The mechanism of action of PT150 involves modulation of the glucocorticoid and androgen receptors that inhibits viral entry and replication by decreasing expression of two main proteins that the virus utilizes for endosomal uptake, TMPRSS2 and ACE2 [8, 35–40]. PT150 also modulates the immune response likely through the glucocorticoid receptor, which is expressed in numerous immune cells including macrophages, T cells, dendritic cells and epithelial cells [41, 42]. PT150 was administered orally to animals infected with SARS-CoV-2, resulting in marked clinical and pathological improvements when compared to the infected vehicle control group. Because Syrian hamsters can propagate human SARS-CoV-2 and show a high degree of sequence homology in both AR and GR (**Fig 4**), it is likely that SARS-CoV-2 infected patients will response similarly to PT150 treatment.

Clinical symptoms of SARS-CoV-2 infection were observed in infected vehicle-only treated animals as early 3DPI. Behavioral and physical changes were more prominent at the 5-day timepoint and included labored breathing, ruffled fur, akinesia and maximal weight loss (5-10%). Weight loss observed 5DPI similar to other studies in Syrian hamsters [43–46]. Animals that received PT150 at clinically relevant low (30mg/kg/day) and high (100mg/Kg/day) doses did not display as severe clinical manifestations and weight loss was minimal at the 5-day timepoint, when compared to untreated control animals (**Fig. 1**). Pathological analysis of infected lung tissue revealed inflammatory cellular infiltration involvement at 3DPI, of which peaked at 5DPI and included mixed inflammatory cell populations resulting in bronchiointerstital pneumonia (**Fig. 2)**. The majority of the parenchymal space at the 5-day timepoint was reduced due to intense inflammatory cell infiltration and showed extravasating cells from pulmonary vessels being recruited to the epithelial layers of the bronchi. The multifocal inflammation began to resolve by 7DPI and was replaced with fibrotic lung tissue and consolidated areas of focal inflammation. Inflammatory pathology within the treated animals was decreased in severity and showed marked improvement within the parenchymal and alveolar spaces. High dose treatment tissue resembled that of control with open alveolar interfaces and non-inflamed vessels.

Digital image analysis of whole-lung scans in the infected vehicle-only group (**Fig 3**) showed extensive hypercellularity and bronchiointerstital pneumonia, with markedly increased hypercellularity in bronchioles, pulmonary arteries and the lung parenchyma. Lung hypercellularity in infected animals increased throughout the course of the 7-day study, similar to the progressive pneumonia experienced by SARS-CoV-2 patients in response to the overwhelming amount of edema and cellular infiltration into the parenchyma of the lungs. Treatment with PT150 markedly reduced hypercellularity at 5 and 7DPI, with the low-dose PT150 group showing reduction at 7DPI. This indicates that PT150 may be regulating cellular infiltration and recruitment into the lung tissue, resulting in reduced pathology and inflammation.

We performed phylogenetic analysis and molecular docking studies to characterize the putative molecular targets by which PT150 could modulate disease progression (**Fig 4**). Sequence concordance values for both AR and GR were very high across species representative of divergent taxonomic groups, including the greater horseshoe bat, sunda pangolin, Syrian golden hamster and human (amongst other species). This underscores not only the likely similar patterns of regulation of AR/GR-dependent genes across species that are required for entry of SARS-CoV-2, such as ACE2 and TMPRSS2, but also the predictive potential for therapeutic modulation of these receptors using PT150, given the sequence similarity between the hamster and human genes. Similarity in these regulatory proteins also highlights the potential for cross-species transmission of SARS-CoV-2 and related β-coronaviruses and the importance of identifying compounds that can modulate signaling of AR and GR to increase host defense.

Molecular docking studies (**Fig 4C,D**) indicated likely interactions with the co-activator site of the ligand binding domain of both receptors (shaded in green), rather than the steroid binding pocket (shaded in magenta in each structure). Thus, PT150 may functions as an allosteric modulator of these receptors, resulting in transcriptional repression of key genes, such as TMRPSS2 and ACE2 (required for viral entry), as well as pro-inflammatory factors. Modeling was conducted using the published crystal structures for AR (PDB: 2PIT) [22] and the glucocorticoid receptor (PDB: 3CLD), solved as a complex co-crystalized with flutacasone [23]. PT150 did not bind the steroid site in any simulation and although the binding site has some flexibility in both receptors, the stereochemistry of PT150 appears to hinder binding at this site. The co-activator peptide binding site most favorably interacted with PT150, suggesting this as a site of allosteric modulation that could inhibit transcriptional activity without directly competing for binding with endogenous corticosteroid ligands. This has been reported for small molecule modulators of other nuclear receptors, such as NR4A2/Nurr1, where molecular docking studies indicated interactions at the co-activator peptide binding site than correlated with transcriptional inhibition of inflammatory gene expression [47]. Studies published with the 1NHZ structure of GR co-crystalized with RU486 suggested potential interactions within the steroid binding pocket of the LBD [48], but the helix that forms part of both the steroid site and the co-activator peptide site was not complete in the 1NHZ structure due to a 9 amino acid deletion, resulting in a distortion of the GR LBD that would be more permissive of interactions within the steroid binding pocket. Thus, PT150 may modulate both AR and GR signaling as an allosteric inhibitor through interactions with the co-activator domain, leading to decreased transcription of target genes.

The doses of PT150 administered in the *in vivo* studies in hamsters are highly comparable to the calculated human doses in the proposed Phase II trial. Previous clinical steady-state exposure data in human trials with 500 mg PT150 would yield a dose of approximately 7 mg/Kg. Compared to both rodent and dog, human calculated exposures are significantly greater for a given dose, based on standard physiologic-based pharmacokinetic models that account for differences in mass, body surface area (BSA), metabolism, half-life, etc. [49]. Using scaling factors based on BSA (33), the doses used in the hamster study translate to a Human Equivalent Dose (HED) of: HED = 30 mg/kg × 5/37 = 4.1 mg/kg; HED = 100 mg/kg × 5/37 = 13.5 mg/kg. Thus, 900 mg/day in patients would represent approximately 12.6 mg/Kg, which is very close to the efficacious dose observed in hamsters treated with PT150.

To further investigate the molecular targets of PT150 *in vivo*, expression of ACE2 was determined in lung in bronchiolar epithelial cells (**Fig 5)**. ACE2 levles in untreated animals remained constant until preaking at 7DPI. Co-localization of SARS-CoV-2 and ACE2 were observed in bronchiolar epithelial cells at 3DPI and 5DPI, supporting that ACE2 is a vital component in viral binding and entry (**Fig 5A-C**). This demonstrates that viral infectivity and replication may transcriptionally regulate ACE2 production in favor of viral propagation and survival. Treatment with 100 mg/Kg PT150 decreased the overall amount of ACE2 in addition to reducing co-localization with SARS-CoV-2. This reduction of ACE2 could be due to inhibition of androgen receptor binding to the promoter of *Ace2*, thereby decreasing transcriptional activation, as reported for other anti-androgens [50]. Expression of TMPRSS2 was also investigated due to the requirement of this cell surface serine protease for processing of the viral S1 spike protein that is necessary for viral entry in complex with ACE2 (**Fig 6**). Protein levels of TMPRSS2 in infected vehicle-only treated animals increased significantly at all timepoints, demonstrating transcriptional induction favoring viral replication. However, the low-dose PT150 treatment decreased TMPRSS2 expression at 5 and 7DPI, whereas high-dose PT150 treatment decreased TMPRSS2 expression at all timepoints, demonstrating a dose-response effect in modulating expression of TMPRSS2 in lung. Decreased production of both TMPRSS2 and ACE2 suggests that PT150 is a transcriptional inhibitor of these genes that likely prevents binding of co-activator proteins to the enhancer regions of TMPRSS2 [51] as well as ACE2 [50]. These data suggest that the anti-viral activity of PT150 is due to direct modulation of host defense that decreases viral entry points.

To determine if PT150 was effective at reducing overall viral replication, infection and immune responses, whole lung sections were analyzed and viral infectivity was determined by assessment of the SARS-CoV-2 nucleocapsid protein (**Fig 7)**. In untreated animals, there was a significant change from control in the percentage of lung area infected with SARS-CoV-2 at 3DPI that decreased by 5DPI and 7DPI, as seen in previously published studies [44, 52, 53]. Viral loads in animals administered PT150 at 100 mg/Kg were not different from control at 3DPI, demonstrating a dramatic reduction in viral attachment, replication and dissemination throughout lung tissue, both in the bronchi and in the parenchyma. Macrophage infiltration per area of tissue in untreated animals was increased at 3DPI and peaked at 5DPI, with resolution occurring at the 7-day timepoint. Animals treated with the high dose of PT150 showed significant decreases in invading macrophage populations at the 3-day timepoint and the low-dose PT150 group showed decreases at 5DPI, demonstrating a marked reductions in the severity of inflammatory infiltration of macrophages.

IL-6 produced by activated macrophages is known to be a critical mediator of lung injury in COVID19 patients [54]. This cytokine is responsible for recruitment and activation of inflammatory cells and is closely associated with the hyperimmune response observed in patients characterized by severe infiltration of macrophages and lymphocytes into lung tissue. IL-6 production was investigated in the bronchiolar cell layer to determine inflammatory recruitment potential during the course of disease (**Fig 8**). Infected animals treated with vehicle showed peak IL-6 production at the 3DPI timepoint, decreasing steadily to the 7-day timepoint. Interestingly, high and low dose PT150 treated groups showed highly reduced levels of IL-6 at all timepoints. This highlights the efficacy of PT150 in reducing innate immune responses concomitant to mitigating the severity of lung pathology in response to SARS-CoV-2.

In conclusion, these data demonstrate disease progression and pathology within the lungs of SARS-CoV-2-infected animals depends upon production of both ACE2 and TMPRSS2, which facilitate viral entry and replication, leading to recruitment of macrophages that initiate a severe innate immune response leading to broncho-interstitial pneuomonia and consolidation of the lung parenchyma. Expression of IL-6 is necessary for recruitment of immune cells to the site of infection, which leads to the ‘cytokine storm’ and decreased prognosis for patients [55, 56]. PT150 treatment interrupts this progression of disease, limiting viral entry, thereby reducing viral loads and decreasing the severity of the immune response to SARS-CoV-2 infection. This may occur through allosteric inhibition of the androgen and glucocorticoid receptors, which decreases protein levels of ACE2 and TMPRSS2 and mitigates excessive immune responses through inhibition of inflammatory gene expression. Importantly, decreased production of IL-6 production by resident immune cells within the lung tissue is likely to correlate with an improve prognosis for patients. The novel mechanism of action of PT150 as both an inhibitor of viral entry and an immunomodulator makes it a strong candidate for therapeutic intervention in the treatment of COVID19 independent of SARS-CoV-2 variants.

## Notes

### Competing Interest Statement

The authors have declared no competing interest.

## References

1. Zhu, N., et al., A Novel Coronavirus from Patients with Pneumonia in China, 2019. N Engl J Med, 2020. 382(8): p. 727–733.

2. Huang, C., et al., Clinical features of patients infected with 2019 novel coronavirus in Wuhan, China. Lancet, 2020. 395(10223): p. 497–506.

3. WHO, Coronavirus dicease (COVID-19)pandemic. 2020.

4. Wiersinga, W.J., et al., Pathophysiology, Transmission, Diagnosis, and Treatment of Coronavirus Disease 2019 (COVID-19): A Review. JAMA, 2020. 324(8): p. 782–793.

5. Fehr, A.R. and S. Perlman, Coronaviruses: an overview of their replication and pathogenesis. Methods Mol Biol, 2015. 1282: p. 1–23.

6. Belouzard, S., V.C. Chu, and G.R. Whittaker, Activation of the SARS coronavirus spike protein via sequential proteolytic cleavage at two distinct sites. Proc Natl Acad Sci U S A, 2009. 106(14): p. 5871–6.

7. Fuentes-Prior, P., Priming of SARS-CoV-2 S protein by several membrane-bound serine proteinases could explain enhanced viral infectivity and systemic COVID-19 infection. J Biol Chem, 2020.

8. Qiao, Y., et al., Targeting transcriptional regulation of SARS-CoV-2 entry factors ACE2 and TMPRSS2. Proc Natl Acad Sci U S A, 2020.

9. Goren, A., et al., Anti-androgens may protect against severe COVID-19 outcomes: results from a prospective cohort study of 77 hospitalized men. J Eur Acad Dermatol Venereol, 2020.

10. Sang, E.R., et al., Epigenetic Evolution of ACE2 and IL-6 Genes: Non-Canonical Interferon-Stimulated Genes Correlate to COVID-19 Susceptibility in Vertebrates. Genes (Basel), 2021. 12(2).

11. Wang, M., et al., Distinct expression of SARS-CoV-2 receptor ACE2 correlates with endotypes of chronic rhinosinusitis with nasal polyps. Allergy, 2020.

12. Russell, C.D., J.E. Millar, and J.K. Baillie, Clinical evidence does not support corticosteroid treatment for 2019-nCoV lung injury. Lancet, 2020. 395(10223): p. 473–475.

13. Saheb Sharif-Askari, N., et al., Effect of Common Medications on the Expression of SARS-CoV-2 Entry Receptors in Kidney Tissue. Clin Transl Sci, 2020. 13(6): p. 1048–1054.

14. Sinha, S., et al., In vitro and in vivo identification of clinically approved drugs that modify ACE2 expression. Mol Syst Biol, 2020. 16(7): p. e9628.

15. Guan, B.J., H.Y. Wu, and D.A. Brian, An optimal cis-replication stem-loop IV in the 5′ untranslated region of the mouse coronavirus genome extends 16 nucleotides into open reading frame 1. J Virol, 2011. 85(11): p. 5593–605.

16. Josset, L., et al., Cell host response to infection with novel human coronavirus EMC predicts potential antivirals and important differences with SARS coronavirus. mBio, 2013. 4(3): p. e00165–13.

17. Glass, C.K. and K. Saijo, Nuclear receptor transrepression pathways that regulate inflammation in macrophages and T cells. Nat Rev Immunol, 2010. 10(5): p. 365–76.

18. Shaked, I., et al., Transcription factor Nr4a1 couples sympathetic and inflammatory cues in CNS-recruited macrophages to limit neuroinflammation. Nat Immunol, 2015. 16(12): p. 1228–34.

19. Peeters, B.W., et al., Glucocorticoid receptor antagonists: new tools to investigate disorders characterized by cortisol hypersecretion. Stress, 2004. 7(4): p. 233–41.

20. Rice, B.A., et al., Repeated subcutaneous administration of PT150 has dose-dependent effects on sign tracking in male Japanese quail. Exp Clin Psychopharmacol, 2019. 27(6): p. 515–521.

21. Theise, N.D., et al., Clinical stage molecule PT150 is a modulator of glucocorticoid and androgen receptors with antiviral activity against SARS-CoV-2. Cell Cycle, 2020. 19(24): p. 3632–3638.

22. Estebanez-Perpina, E., et al., A surface on the androgen receptor that allosterically regulates coactivator binding. Proc Natl Acad Sci U S A, 2007. 104(41): p. 16074–9.

23. Biggadike, K., et al., X-ray crystal structure of the novel enhanced-affinity glucocorticoid agonist fluticasone furoate in the glucocorticoid receptor-ligand binding domain. J Med Chem, 2008. 51(12): p. 3349–52.

24. Friesner, R.A., et al., Glide: a new approach for rapid, accurate docking and scoring. 1. Method and assessment of docking accuracy. J Med Chem, 2004. 47(7): p. 1739–49.

25. Friesner, R.A., et al., Extra precision glide: docking and scoring incorporating a model of hydrophobic enclosure for protein-ligand complexes. J Med Chem, 2006. 49(21): p. 6177–96.

26. Halgren, T.A., Identifying and characterizing binding sites and assessing druggability. J Chem Inf Model, 2009. 49(2): p. 377–89.

27. Beckstein, O., A. Fourrier, and B.I. Iorga, Prediction of hydration free energies for the SAMPL4 diverse set of compounds using molecular dynamics simulations with the OPLS-AA force field. J Comput Aided Mol Des, 2014. 28(3): p. 265–76.

28. Chan, J.F., et al., Simulation of the Clinical and Pathological Manifestations of Coronavirus Disease 2019 (COVID-19) in a Golden Syrian Hamster Model: Implications for Disease Pathogenesis and Transmissibility. Clin Infect Dis, 2020. 71(9): p. 2428–2446.

29. Imai, M., et al., Syrian hamsters as a small animal model for SARS-CoV-2 infection and countermeasure development. Proc Natl Acad Sci U S A, 2020. 117(28): p. 16587–16595.

30. Young, M.J., C.D. Clyne, and K.E. Chapman, Endocrine aspects of ACE2 regulation: RAAS, steroid hormones and SARS-CoV-2. J Endocrinol, 2020. 247(2): p. R45–R62.

31. Cascella, M., et al., Features, Evaluation, and Treatment of Coronavirus, in StatPearls. 2020: Treasure Island (FL).

32. Asselah, T., et al., COVID-19: Discovery, diagnostics and drug development. J Hepatol, 2021. 74(1): p. 168–184.

33. Campos, D.M.O., et al., SARS-CoV-2 virus infection: Targets and antiviral pharmacological strategies. J Evid Based Med, 2020. 13(4): p. 255–260.

34. Li, H., et al., Impact of corticosteroid therapy on outcomes of persons with SARS-CoV-2, SARS-CoV, or MERS-CoV infection: a systematic review and meta-analysis. Leukemia, 2020. 34(6): p. 1503–1511.

35. Hoffmann, M., et al., SARS-CoV-2 Cell Entry Depends on ACE2 and TMPRSS2 and Is Blocked by a Clinically Proven Protease Inhibitor. Cell, 2020. 181(2): p. 271–280 e8.

36. Sungnak, W., et al., SARS-CoV-2 entry factors are highly expressed in nasal epithelial cells together with innate immune genes. Nat Med, 2020. 26(5): p. 681–687.

37. Lukassen, S., et al., SARS-CoV-2 receptor ACE2 and TMPRSS2 are primarily expressed in bronchial transient secretory cells. EMBO J, 2020. 39(10): p. e105114.

38. Ragia, G. and V.G. Manolopoulos, Inhibition of SARS-CoV-2 entry through the ACE2/TMPRSS2 pathway: a promising approach for uncovering early COVID-19 drug therapies. Eur J Clin Pharmacol, 2020. 76(12): p. 1623–1630.

39. Bourgonje, A.R., et al., Angiotensin-converting enzyme 2 (ACE2), SARS-CoV-2 and the pathophysiology of coronavirus disease 2019 (COVID-19). J Pathol, 2020. 251(3): p. 228–248.

40. Ali, A. and R. Vijayan, Dynamics of the ACE2-SARS-CoV-2/SARS-CoV spike protein interface reveal unique mechanisms. Sci Rep, 2020. 10(1): p. 14214.

41. Baschant, U. and J. Tuckermann, The role of the glucocorticoid receptor in inflammation and immunity. J Steroid Biochem Mol Biol, 2010. 120(2–3): p. 69–75.

42. Necela, B.M. and J.A. Cidlowski, Mechanisms of glucocorticoid receptor action in noninflammatory and inflammatory cells. Proc Am Thorac Soc, 2004. 1(3): p. 239–46.

43. Sia, S.F., et al., Pathogenesis and transmission of SARS-CoV-2 in golden hamsters. Nature, 2020. 583(7818): p. 834–838.

44. Rosenke, K., et al., Defining the Syrian hamster as a highly susceptible preclinical model for SARS-CoV-2 infection. bioRxiv, 2020.

45. Lee, A.C., et al., Oral SARS-CoV-2 Inoculation Establishes Subclinical Respiratory Infection with Virus Shedding in Golden Syrian Hamsters. Cell Rep Med, 2020. 1(7): p. 100121.

46. Brocato, R.L., et al., Disruption of Adaptive Immunity Enhances Disease in SARS-CoV-2 Infected Syrian Hamsters. J Virol, 2020.

47. Popichak, K.A., et al., Compensatory Expression of Nur77 and Nurr1 Regulates NF-kappaB-Dependent Inflammatory Signaling in Astrocytes. Mol Pharmacol, 2018. 94(4): p. 1174–1186.

48. Kauppi, B., et al., The three-dimensional structures of antagonistic and agonistic forms of the glucocorticoid receptor ligand-binding domain: RU-486 induces a transconformation that leads to active antagonism. J Biol Chem, 2003. 278(25): p. 22748–54.

49. Reagan-Shaw, S., M. Nihal, and N. Ahmad, Dose translation from animal to human studies revisited. FASEB J, 2008. 22(3): p. 659–61.

50. Deng, Q., et al., Targeting androgen receptor regulation of TMPRSS2 and ACE2 as a therapeutic strategy to combat CoVID-19 2021.

51. Martinez-Ariza, G. and C. Hulme, Recent advances in allosteric androgen receptor inhibitors for the potential treatment of castration-resistant prostate cancer. Pharm Pat Anal, 2015. 4(5): p. 387–402.

52. Osterrieder, N., et al., Age-Dependent Progression of SARS-CoV-2 Infection in Syrian Hamsters. Viruses, 2020. 12(7).

53. Roberts, A., et al., Severe acute respiratory syndrome coronavirus infection of golden Syrian hamsters. J Virol, 2005. 79(1): p. 503–11.

54. Stasi, C., et al., Treatment for COVID-19: An overview. Eur J Pharmacol, 2020. 889: p. 173644.

55. Hojyo, S., et al., How COVID-19 induces cytokine storm with high mortality. Inflamm Regen, 2020. 40: p. 37.

56. Tang, L., et al., Controlling Cytokine Storm Is Vital in COVID-19. Front Immunol, 2020. 11: p. 570993.

